# Amilorides inhibit SARS-CoV-2 replication in vitro by targeting RNA structures

**DOI:** 10.1101/2020.12.05.409821

**Authors:** Martina Zafferani, Christina Haddad, Le Luo, Jesse Davila-Calderon, Liang Yuan-Chiu, Christian Shema Mugisha, Adeline G. Monaghan, Andrew A. Kennedy, Joseph D. Yesselman, Robert R. Gifford, Andrew W. Tai, Sebla B. Kutluay, Mei-Ling Li, Gary Brewer, Blanton S. Tolbert, Amanda E. Hargrove

**Author notes:** Co-first authors (Martina Zafferani, Christina Haddad). Co-second authors (Le Luo, Jesse Davila-Calderon, Liang Yuan-Chiu).

## Abstract

The SARS-CoV-2 pandemic, and the likelihood of future coronavirus pandemics, has rendered our understanding of coronavirus biology more essential than ever. Small molecule chemical probes offer to both reveal novel aspects of virus replication and to serve as leads for antiviral therapeutic development. The RNA-biased amiloride scaffold was recently tuned to target a viral RNA structure critical for translation in enterovirus 71, ultimately uncovering a novel mechanism to modulate positive-sense RNA viral translation and replication. Analysis of CoV RNA genomes reveal many conserved RNA structures in the 5’-UTR and proximal region critical for viral translation and replication, including several containing bulge-like secondary structures suitable for small molecule targeting. Following phylogenetic conservation analysis of this region, we screened an amiloride-based small molecule library against a less virulent human coronavirus, OC43, to identify lead ligands. Amilorides inhibited OC43 replication as seen in viral plaque assays. Select amilorides also potently inhibited replication competent SARS-CoV-2 as evident in the decreased levels of cell free virions in cell culture supernatants of treated cells. Reporter screens confirmed the importance of RNA structures in the 5’-end of the viral genome for small molecule activity. Finally, NMR chemical shift perturbation studies of the first six stem loops of the 5’-end revealed specific amiloride interactions with stem loops 4, 5a, and 6, all of which contain bulge like structures and were predicted to be strongly bound by the lead amilorides in retrospective docking studies. Taken together, the use of multiple orthogonal approaches allowed us to identify the first small molecules aimed at targeting RNA structures within the 5’-UTR and proximal region of the CoV genome. These molecules will serve as chemical probes to further understand CoV RNA biology and can pave the way for the development of specific CoV RNA-targeted antivirals.

## Introduction

A new strain of severe acute respiratory syndrome coronavirus 2 (SARS-CoV-2) was reported in November 2019 from Wuhan, China. SARS-CoV-2 is the etiological agent of the COVID-19 respiratory disease, the largest scale respiratory virus pandemic the world has witnessed since the 1918 Spanish flu^1^ and that has claimed more than 1.4 million lives worldwide.^2^ Coronaviruses (CoV family) generally cause mild flu-like symptoms^3^ in humans but have caused two smaller scale pandemics in the last two decades: SARS-CoV (2003) and MERS (2012)^4^. Recent phylogenetic mapping traced all human coronaviruses to animal origins.^5^ While the middle zoonotic carrier of the virus between the animal of origin and humans seem to vary between CoVs, the chronological surfacing of human CoV pandemics seems to follow a dangerous trend of increasing lethality of each pandemic, thereby underscoring the need to a better understanding and targeting of the current and future CoV etiologic agents.

After more than a year since the first cases of SARS-CoV-2 human infection, this virus is expected to remain a global threat until vaccines are available and adopted worldwide. While recent treatments have been approved for use within hospital settings,^6^ there are no known FDA approved cures for the infection. Current candidates for treatment have limited approval for emergency use in severe COVID-19 cases. Remdesivir, for example, is an RNA-dependent RNA polymerase inhibitor (RdRp) initially developed during the Ebola outbreak and revisited at the start of the pandemic.^7^ The compassionate use of the candidate antiviral across many countries reported mixed results, with overall faster recovery time from the virus but no difference in mortality rates.^8^ While more randomized trials are needed for a final verdict on the efficacy of Remdesivir in critical patients, its stereospecific multi-step synthetic process highlights the need for new, scalable and more efficacious antivirals. Baricitinib has been recently approved for emergency use for COVID-19 treatment in conjunction with Remdesivir. Also known as Olumiant, this small molecule was approved in 2018 as treatment for moderate to severe rheumatoid arthritis.^9^ It is proposed that the anti-inflammatory effects of the drug help in decreasing inflammatory cascades associated with COVID-19. While promising, Baricitinib has yet to receive approval as a stand-alone treatment and, so far, has been shown to improve recovery time by one day when compared to Remdesivir alone treatment.^6^ Overall, SARS-CoV-2 is not the first and, most likely, will not be the last CoV pandemic. The current limited tools and lack of novel cures underscore the need for a new approach in developing antivirals that would not only provide novel routes to combat the current pandemic but also provide invaluable information on targetable structures that can aid in the prevention of and fight against future CoV outbreaks.

Several steps in the coronavirus replication cycle offer potential therapeutic targets for viral inhibition (**Figure 1**). Coronaviruses are enveloped positive single-stranded RNA genomes of approximately 30 kilobases, making them the largest genomes of RNA viruses.^4^ SARS-CoV-2 infects human cells by engagement of the ACE2 receptor by the viral spike (S) protein followed by membrane fusion at the plasma membrane or endosomal membranes depending on the availability of host cell proteases that cleave and prime S for entry. Fusion results in the release of its positive sense genome and associated proteins in the host cell cytosol. The genome is translated into two large polyproteins, 1a and 1b, which are then processed into individual proteins by the viral protease. Synthesis of full length negative strand RNA by products of 1a/1b creates a template to synthesize multiple positive strand copies encapsidated by the viral nucleoprotein into virions.^4^ The negative RNA strand also serves as a template for the synthesis of shorter sub-genomic RNAs (sgRNA) that include the essential structural proteins and thus also constitute an attractive therapeutic target.^10^

**Figure 1.**
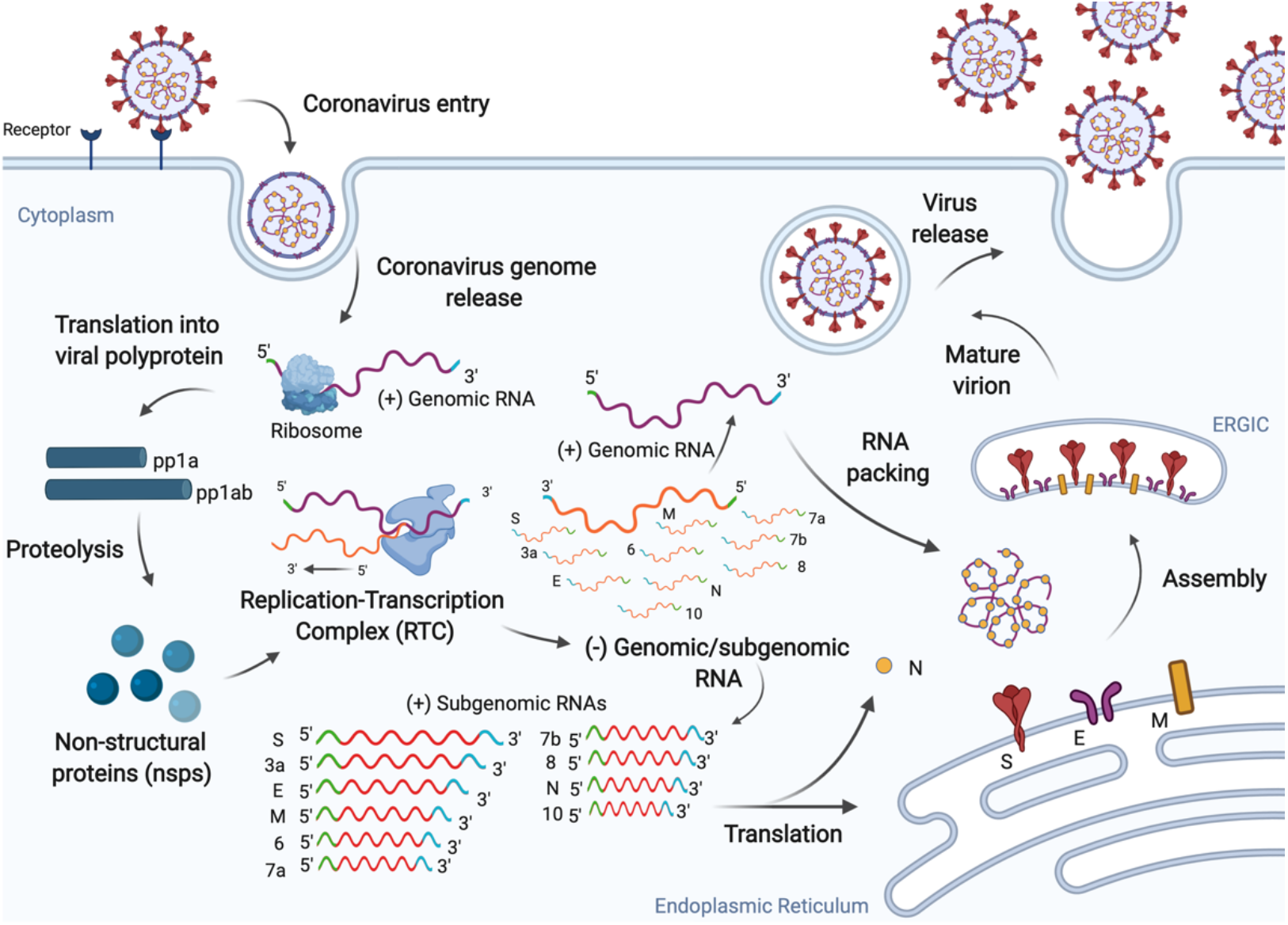
SARS-CoV-2 replication cycle. The virus enters human cells via endocytosis by binding the ACE2 receptor and releasing its positive-sense RNA genome. The virus exploits the host machinery to facilitate efficient viral replication, which ultimately leads to progression of infection.^11^

CoV antivirals to date have been developed to target viral proteins, including to prevent endocytosis, assembly of viral protein for export, and condensation of viral genome for packaging.^12^ While this protein-centric approach has proven successful in a few cases, the sequence and structural conservation of RNA structural motifs pose an attractive complementary target^13^ for small molecule antiviral development, a strategy that has shown promise against a plethora of viruses.^14^ Recently published data on genome-wide secondary structure of the virus obtained by *in vitro* and *in vivo* SHAPE ^15-17^ and DMS^17-19^ probing on SARS-CoV-2 infected Vero E6 cells, as well as NMR characterization,^19^ recapitulates the computationally predicted stem loops (SL) at the 5’ region of the genome (**Figure 2**) as well other relevant frameshifting and replication-related structures. Conserved elements at the 5’ and 3’-ends have been identified across many members of the coronavirus family and function as *cis* acting elements regulating viral replication.^20^ Specifically, studies on murine and bovine coronaviruses showed that phylogenetically conserved stem loops in the 5’-UTR are capable of long-range RNA-RNA and protein-RNA interactions responsible for optimal viral replication.^21,22^ More recently, studies aimed at uncovering the pathway that leads to viral protein synthesis via host cell translation machinery revealed that the presence of the full length 5’-UTR of SARS-CoV-2 leads to a five-fold increase in translation of viral proteins. This preliminary data corroborates the importance of the 5’-UTR region, and the structures within, for efficient viral translation and provides context for the viral hijacking of the host cell translational machinery.^23^

**Figure 2.**
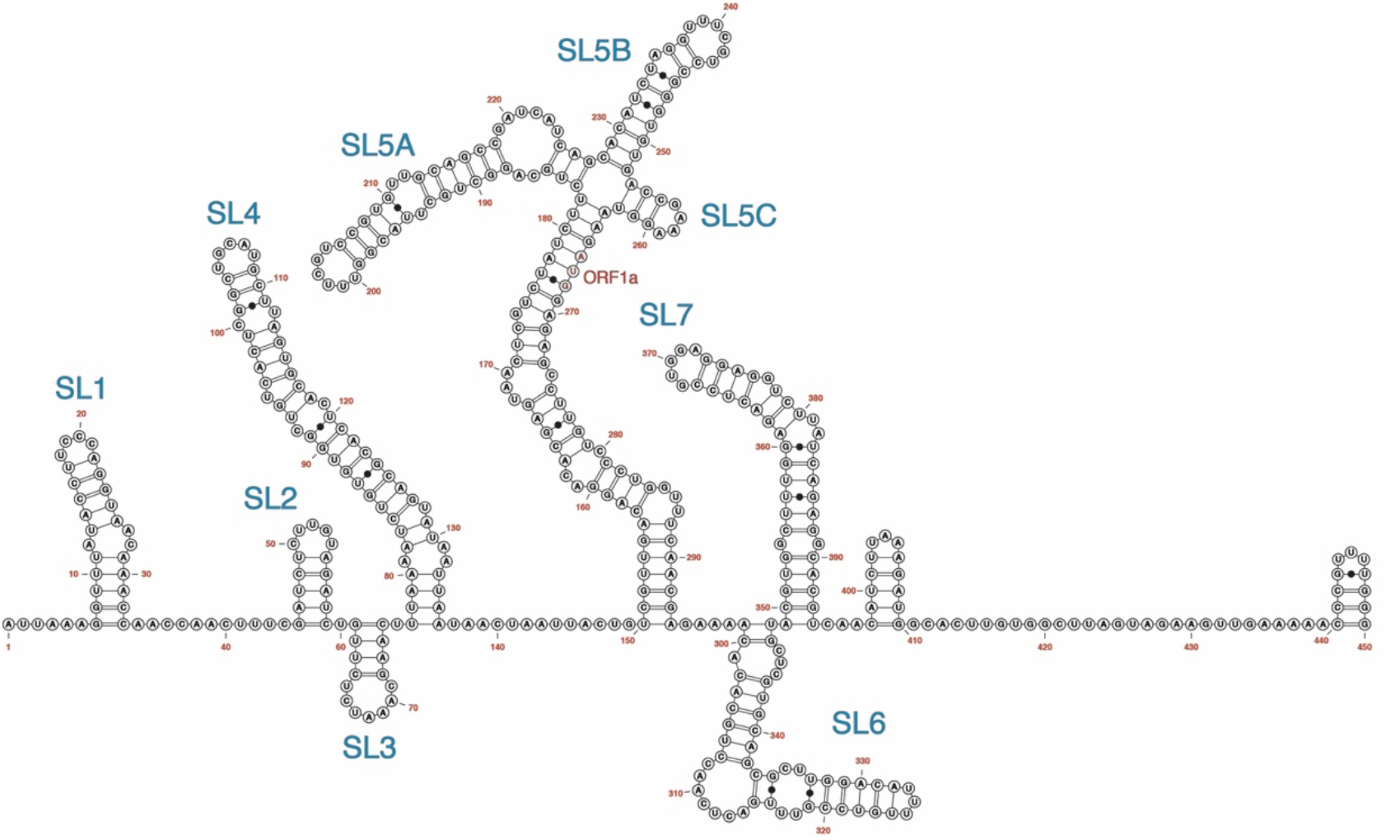
Secondary structure of the 5’-end (450 nt) of the SARS-CoV-2 RNA genome as determined by DMS chemical probing.^19^

Drug-like small molecules offer the ability to develop chemical probes that reveal function and to design bioavailable clinical candidates for treatment. While RNA targeting has lagged behind protein targeting, recent successes in both the laboratory and the clinic support its potential role. The first US FDA approved small molecule targeting RNA other than the ribosome was approved for treatment of spinal muscular atrophy in August of 2020.^24^ Effective small molecule targeting in the laboratory has also been observed for a plethora of disease relevant RNAs, including viral RNAs. Specifically, small molecules targeting structures within the 5’-UTR region have shown antiviral activity for a number of positive-sense RNA viruses such as HCV^25^, FMDV^26^, and EV71.^27^ At the same time, recent studies have begun evaluating the potential of small molecules against the frameshifting elements of SARS-CoV-2 RNA in the coding region. These recent studies include evaluation of known SARS-CoV pseudoknot binders^28^ as well as development of a small molecule binder to the attenuator hairpin preceding the pseudoknot.^29^ Coupling of the latter to the known ribonuclease targeting chimera (RIBOTAC) technology results in recruitment of cellular ribonucleases leading to viral genomic RNA degradation. While the anti-replicative activity of these molecules has not been published, these preliminary results, combined with known RNA-targeted antivirals for other positive sense RNA viruses, stand as proof-of concept of the targetability of SARS-CoV-2 RNA motifs.

Successful efforts to target viral 5’-UTR structures have often leveraged the synthetic tuning of a known RNA binding scaffold.^25, 27, 30^ We recently employed this approach to develop RNA-targeted antivirals based on dimethylamiloride (DMA) that target the internal ribosomal entry site (IRES) region in the 5’-UTR of EV71. The DMA scaffold had been previously reported to be poised for tuning for specific RNA constructs via a facile three-step synthesis.^31^ Further investigation and functionalization of the scaffold resulted in a bioactive antiviral analogue that formed a repressive ternary complex with IRES stem loop II RNA and the human AUF1 protein, ultimately inhibiting translation and compromising viral replication. The successes in DMA exploration highlight the scaffold’s potential for tuning as well as broad applicability as an antiviral scaffold.^31^ Herein we report DMA analogues that show promising antiviral properties by reducing SARS-CoV-2 viral titer in a dose-dependent manner in infected Vero E6 cells. In addition, dual luciferase reporter assays confirmed the antiviral activity of the small molecules to be dependent on the 5’-UTR and proximal region of the SARS-CoV-2 genome. Investigation of possible conserved RNA binding sites of the lead small molecules revealed putative bulge-like binding sites in stem loops 4 and 5a, located in the 5’-UTR of the SARS-CoV-2 genome, as well as in the adjacent stem loop 6 located in ORF1a, supporting both the targetability of 5’-region stem loops and the combination of interdisciplinary methods used here as a promising approach to novel anti-SARS-CoV-2 small molecules.

## Results

### I. Phylogenetic conservation of RNA structural elements

As the functional significance of the 5’-end stem loops (SLs) is still being elucidated, we examined sequence conservation in the 5’-end region across the Betacoronavirus genus as a preliminary approach to assessing the suitability of the known structures in this region as therapeutic targets. We constructed a multiple sequence alignment spanning the 5’UTR and the adjacent region including representatives of all five betacoronavirus subgenera (**Figure 3**). Alignments disclosed the highest degree of conservation in the region encoding SLs 2-3, both of which are relatively short and contain stretches of 5-6 nucleotides that are 100% conserved. By contrast, SL1 and SLs 4-6 are less conserved, but notably, four of the five SLs that span the 5’UTR contain stretches of relatively highly conserved nucleotide sequence (i.e. 70-100% conservation across the genus). The position of these SL’s within or adjacent to the 5’UTR (which is relatively conserved in length across all betacoronaviruses) indicates these conserved regions are likely to represent homologous nucleotides and suggests they are to some degree functionally equivalent.

**Figure 3.**
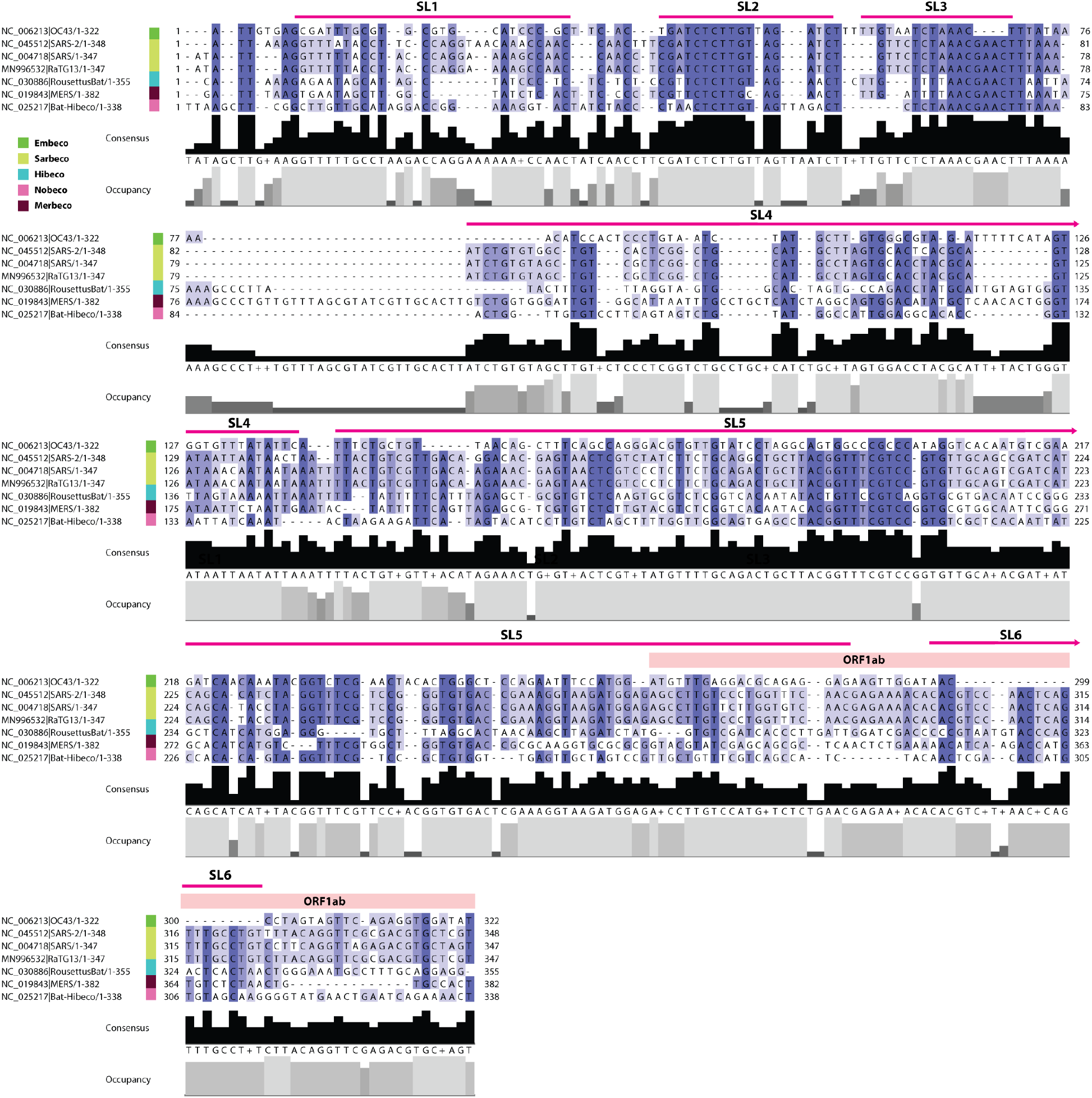
Conservation of 5’-end sequences encoding RNA structures in betacoronavirus genomes. Multiple sequence alignment showing percentage identity and sequence coverage within the 5’ untranslated and adjacent region of representative species within genus Betacoronavirus. Stem loop (SL) structures 2 and 3 are highly conserved, while other stem loops show moderate conservation. Viral subgenus is indicated as shown in the key. Sequence numbering is standardised to the genome sequence start position of each taxon. Taxon labels show GenBank accession numbers and abbreviated virus names. Abbreviations: OC43=Human coronavirus OC43; SARS=Severe acute respiratory syndrome coronavirus; RousettusBat=Rousettus bat coronavirus MERS=Middle East respiratory syndrome-related coronavirus; Bat-Hibeco=Bat Hp-betacoronavirus.

### II. DMAs inhibits human coronavirus OC43 virus replication in a dose-dependent manner

We pursued targeting of the 5’-end stem loop structures with the DMA scaffold, which has been previously reported as an RNA binding scaffold that can be successfully optimized for selectivity for distinct RNA elements.^31^ To quickly assess potential CoV antiviral activity, human OC43 betacoronavirus was used due to its lower virulence and thus suitability for use in standard cell culture facilities.^3^ Vero E6 cells were infected with human coronavirus OC43 at an MOI=1. After a one-hour adsorption, a panel of twenty-three DMAs at 50 μM or 100 µM were added to the cells and incubated for 24 hours at 33°C. Virus titers were determined by plaque formation on Vero E6 cells, and DMA-132, -135, and -155 (**Figure 4**) reduced virus titer by ∼ 1,000-fold at 100 µM concentration **(Figure S1)**. The same experiments were repeated three times with DMA-132, -135, and -155 and shown to be reproducible **(Figure S2)**. The results further suggest DMA-132, - 135, and -155 reduce virus titer in a dose-dependent manner. The parent scaffold, dimethylamiloride (DMA-1), demonstrated no activity and is used as an inactive control moving forward.

**Figure 4.**
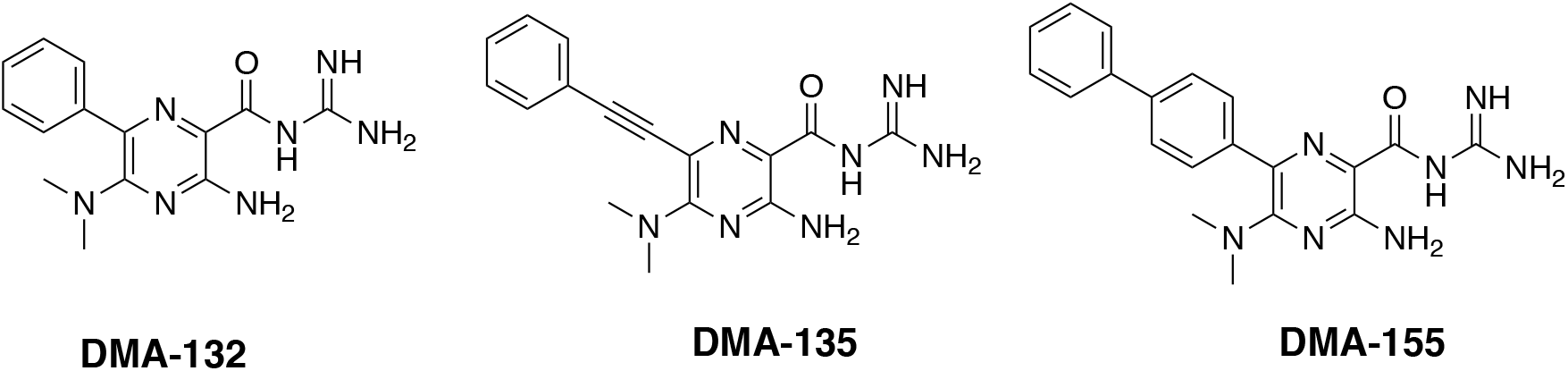
Chemical structure of the three lead molecules derived from the focused library screen against OC43 infected Vero E6 cell screening.

### III. Antiviral potency of DMAs against SARS-CoV-2

To determine the antiviral activity and potency of the lead small molecules against SARS-CoV-2 we utilized a simplified Q-RT-PCR assay to monitor SARS-CoV-2 viral RNA levels in supernatants of infected Vero E6 cells.^32^ Similar to Remdesivir, DMA-135 and 155 led to a dose dependent 10-30-fold decrease in cell-free viral RNA levels within 24 hours of infection with an approximate IC_50_ of 10 and 16 μM, respectively (**Figure 5**).

**Figure 5.**
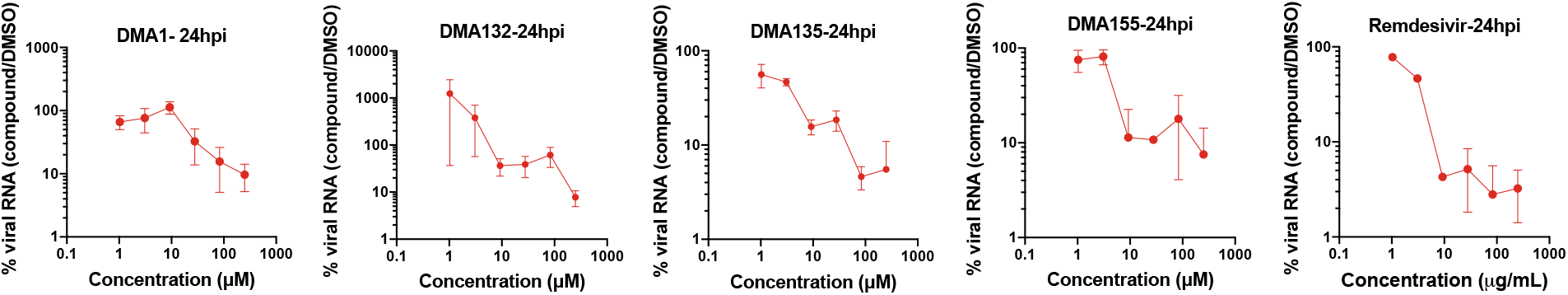
Vero E6 cells were infected with SARS-CoV-2 at MOI: 0.1 i.u./cell and the indicated DMA compounds were added on cells following virus adsorption for 1 hr. Cell culture supernatants were collected and analyzed by a Q-RT-PCR assay using N-specific primers. Data show the relative percentage of viral RNA in DMA-treated samples compared to mock-treated samples. Data are from two independent experiments and error bars show the range. Data for 48- and 72-hour treatment times can be found in Figure S4.

Antiviral activity of the two most active DMA leads (DMA-132 and -135) was confirmed using Vero E6 cells infected with wild-type SARS-CoV-2; DMA-155 was not tested. Small molecule treatment was performed at 10 μM and 50 μM, respectively. DMA-132 and 135 showed significant dose-dependent reduction in viral titer compared to DMSO, as measured by median tissue culture infectious dose (TCID_50_) assay, without measurable effect on cellular viability as measured by ATP content (**Figure 6**). Notably, DMA-132 and 135 have improved activity when compared to DMA-01, the parent scaffold, thereby corroborating the potential for synthetic tunability of DMAs for SARS-CoV-2 targeting.

**Figure 6.**
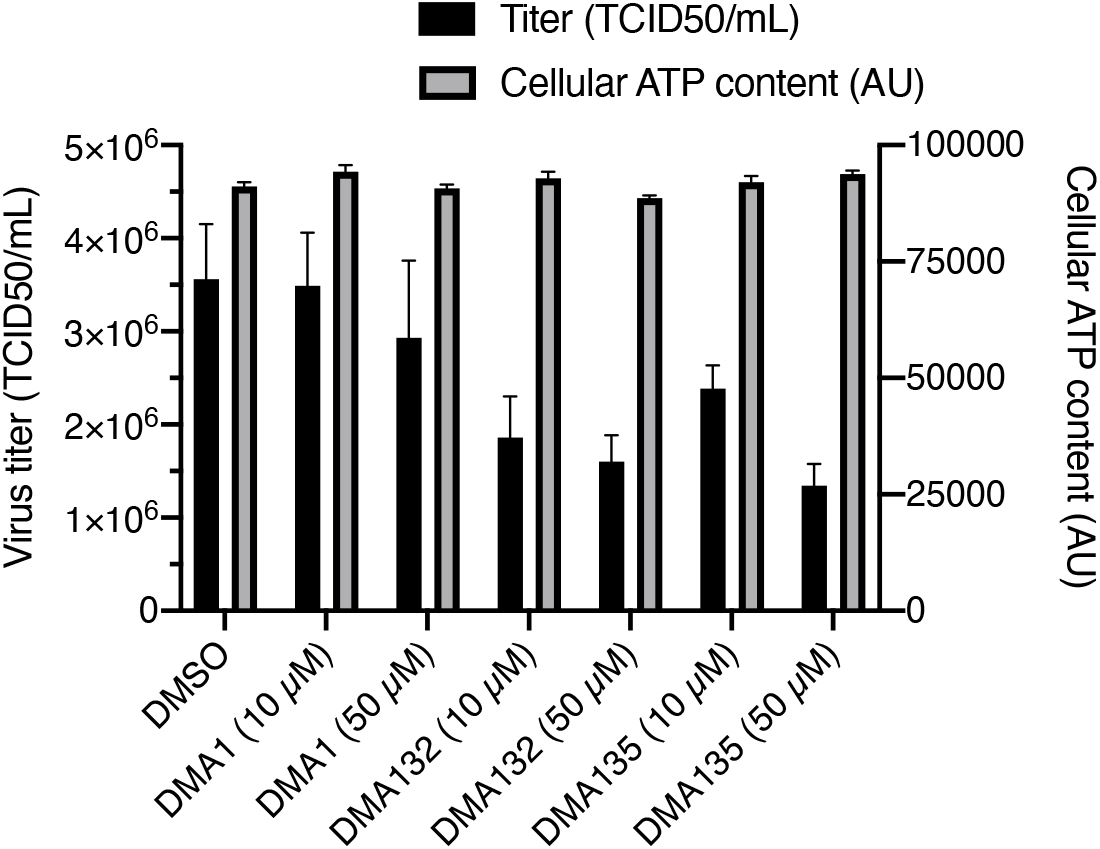
Inhibition of SARS-CoV-2 infection by DMA leads without significant cellular toxicity: VeroE6 cells were infected with SARS-CoV-2 (MOI=0.1) in the presence of DMSO or the indicated compounds for 3 days followed by TCID50 measurement of viral titer in culture supernatant (black bars). Results were generated with two independent experiment, each with two replicates. Cellular viability was assayed by measuring cellular ATP content in uninfected VeroE6 cells after 3 days of treatment with the indicated compounds (grey bars).

DMA-132, 135, and 155 were tested for longer term (96 hour) toxicity in Vero E6 cells. Small molecules do not significantly reduce cell viability < 10 μM, supporting a potential therapeutic window, though more extensive cytotoxicity studies are warranted. In particular, **Figure S3** shows the CC_50_ of DMA-132 and -135 in Vero E6 cells were > 100 µM. CC_50_ of DMA-155 was about 90 µM.

### IV. Investigation of small molecule activity against CoV-2-luciferase reporter gene expression

To assess the effect of DMAs on reporter gene expression directed by SARS-CoV-2 sequence elements, a reporter plasmid, pCoV2-5′UTR-FLuc-3’UTR was used as template for *in vitro* synthesis of CoV-2-5′UTR-FLuc-3’UTR RNA (**Figure 7A**). This plasmid contains the 5’-end 805 nucleotide (nt) segment from the virus genome and the 3’UTR. Thus, the 805 nt segment spans the genomic RNA 5’UTR (SL 1-5) and ORF1a encoding a portion of nsp1 (including SL 6-8) fused in-frame with the firefly luciferase ORF. Plasmid pRL was used as template for the synthesis of control Renilla luciferase reporter RNA lacking SARS-Cov-2 sequences. The RNAs were co-transfected and various concentrations of DMAs were added with the transfections. Two days after transfection, RLuc and FLuc activities were measured using a dual-luciferase reporter assay. As shown in **Figure 7A**, addition of 10 µM DMA-132 or -135 reduced FLuc activity, which is under the control of SARS-CoV-2 5’-end and 3’UTR, by approximately 50%. Addition of 10 µM DMA-155 reduced FLuc activity by ∼ 90%. FLuc activity was reduced 90% with 100 µM of DMA-132, -135 or -155. Activities of the control RLuc remained relatively constant for all DMAs across all concentrations tested; this control indicates that decreases in FLuc were not simply due to any putative cytotoxic effects. To test whether the CoV-2 3’UTR contributes to DMA-mediated translational repression, we repeated the experiment using a FLuc reporter, CoV-2-5’UTR-FLuc RNA, in which the CoV-2 3’UTR was replaced with vector-encoded sequence; thus, the CoV-2 5’-ends of both FLuc reporter RNAs are the same (**Figure 7B**). The effects of DMAs on translational repression were virtually identical to those in **Figure 7A**. Importantly, these results clearly demonstrate that DMA-dependent suppression of SARS-CoV-2 luciferase reporter activity requires only 5’-end sequences of the virus.

**Figure 7.**
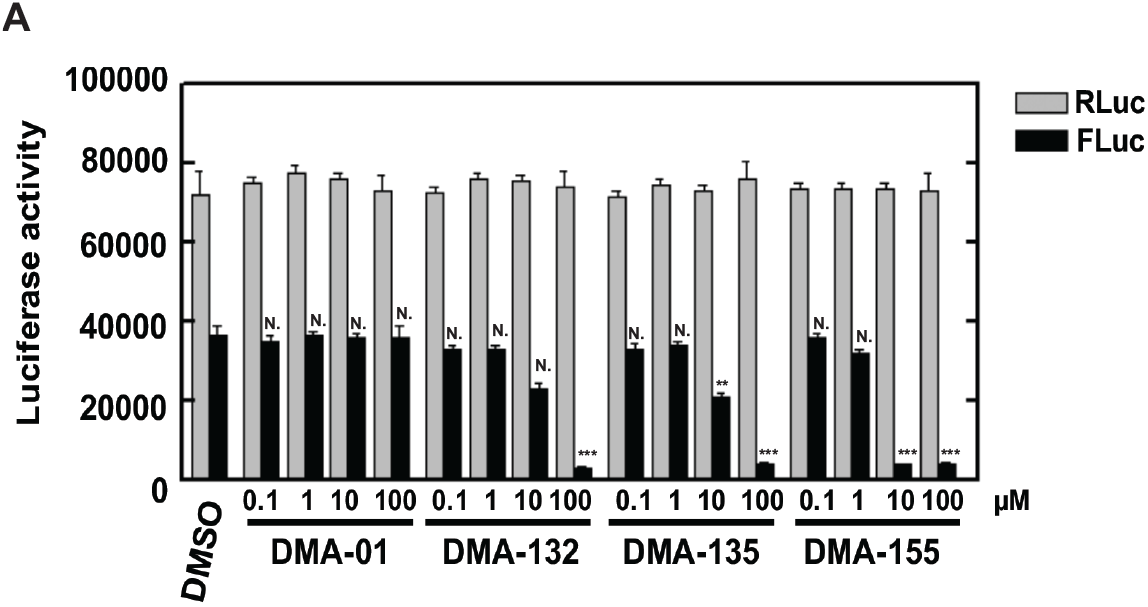
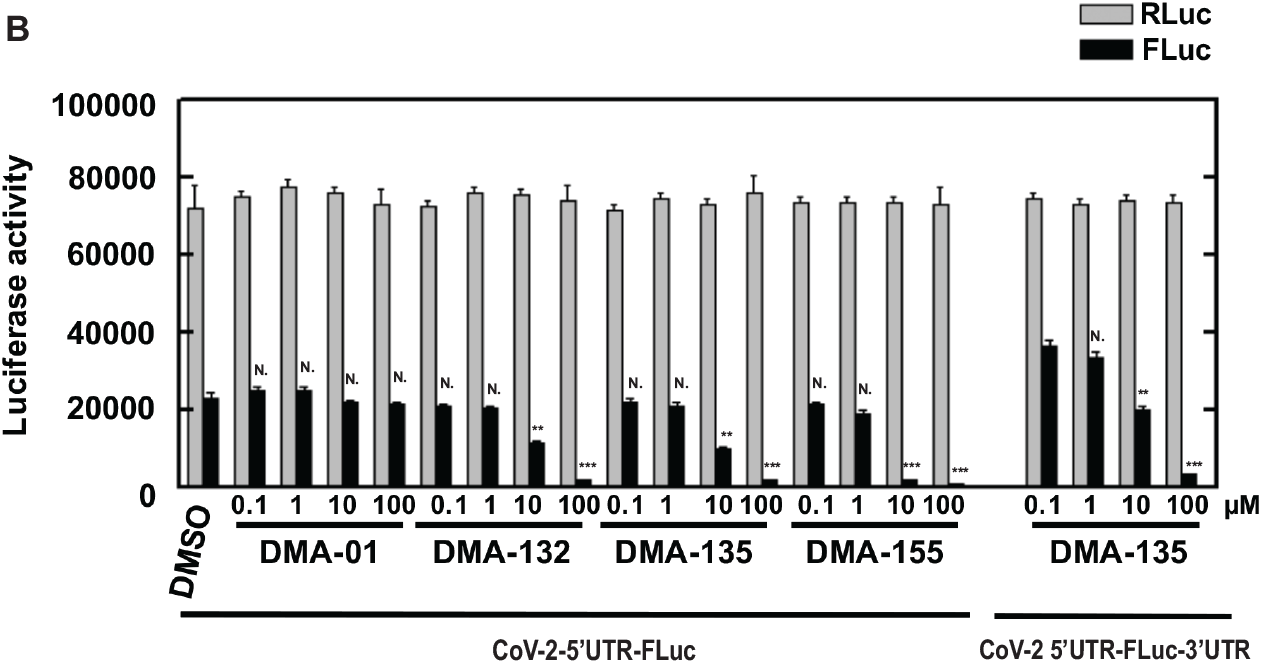
Vero E6 cells were co-transfected with (**A**) CoV-2-5′UTR-FLuc-3’UTR or (**B**) Cov-2-5’UTR-FLuc and control RLuc RNAs and cultured with various concentrations of DMAs. Luciferase activity was measured 2 days later. Mean values and standard deviations from three independent experiments are shown in the bar graphs. \*\*\**P* < 0.001; \*\**P* < 0.01; N., not significant relative to the DMSO control, except for the 5’UTR-FLuc-3’UTR comparison in panel B, which is relative to DMA-135 at 0.1 µM.

### V. NMR Profiling of 5’-Stem loop-DMA interactions

Towards understanding potential mechanisms by which the DMAs inhibit SARS-CoV-2 replication, we carried out single-point ^13^C-^1^H TROSY HSQC titrations of DMA-132, -135 and -155 into stem loops 1-6 (SL1-6) of the 5’-region. We reasoned that the DMAs selectively bind the 5’-region stem loops at non-canonical structural elements such as bulge, internal or apical loops. Therefore, we *in vitro* transcribed isolated SL domains using ^13^C/^15^N rNTPs that would maximize NMR signal detection for the non-canonical elements over the base paired regions (**Figure S5**). For example, SL1 was prepared as a C(^13^C/^15^N)-selectively labeled construct because cytosines are the most abundant nucleobases within its apical and internal loops. Using this strategy, we are able to efficiently profile each 5’-region SL to determine if the DMAs bind with reasonable affinity and specificity. **Figure S5** summarizes the effects that the addition of excess (5:1) DMAs have on the NMR spectra of each major SL domain. First, we observed that the DMAs induced differential chemical shift perturbations (CSPs) for each SL with the largest CSPs observed in spectra recorded for SLs 4, 5a and 6 (**Figure 8**). Each of these 5’-domains contain large internal loops. Second, only a subset of the ^13^C-^1^H correlation peaks is perturbed in the spectra providing evidence that the DMAs interact through specific surfaces of the 5’-region. Third, the extent of the CSPs are also differential with some DMAs inducing shifts of the correlation peaks to new positions within the spectra, and others inducing complete broadening of the NMR signals. These variable CSPs reveal that the DMAs interact with different binding affinities and binding modes. Of note, none of the DMAs caused significant changes to spectra recorded on SL2, which contains a 5-nt CUUGU apical loop. Taken together, the single-point ^13^C-^1^H TROSY HSQC titrations provide compelling evidence that the DMAs make specific interactions with the SARS-CoV-2 5’-region via surfaces composed of non-canonical structural elements.

**Figure 8.**
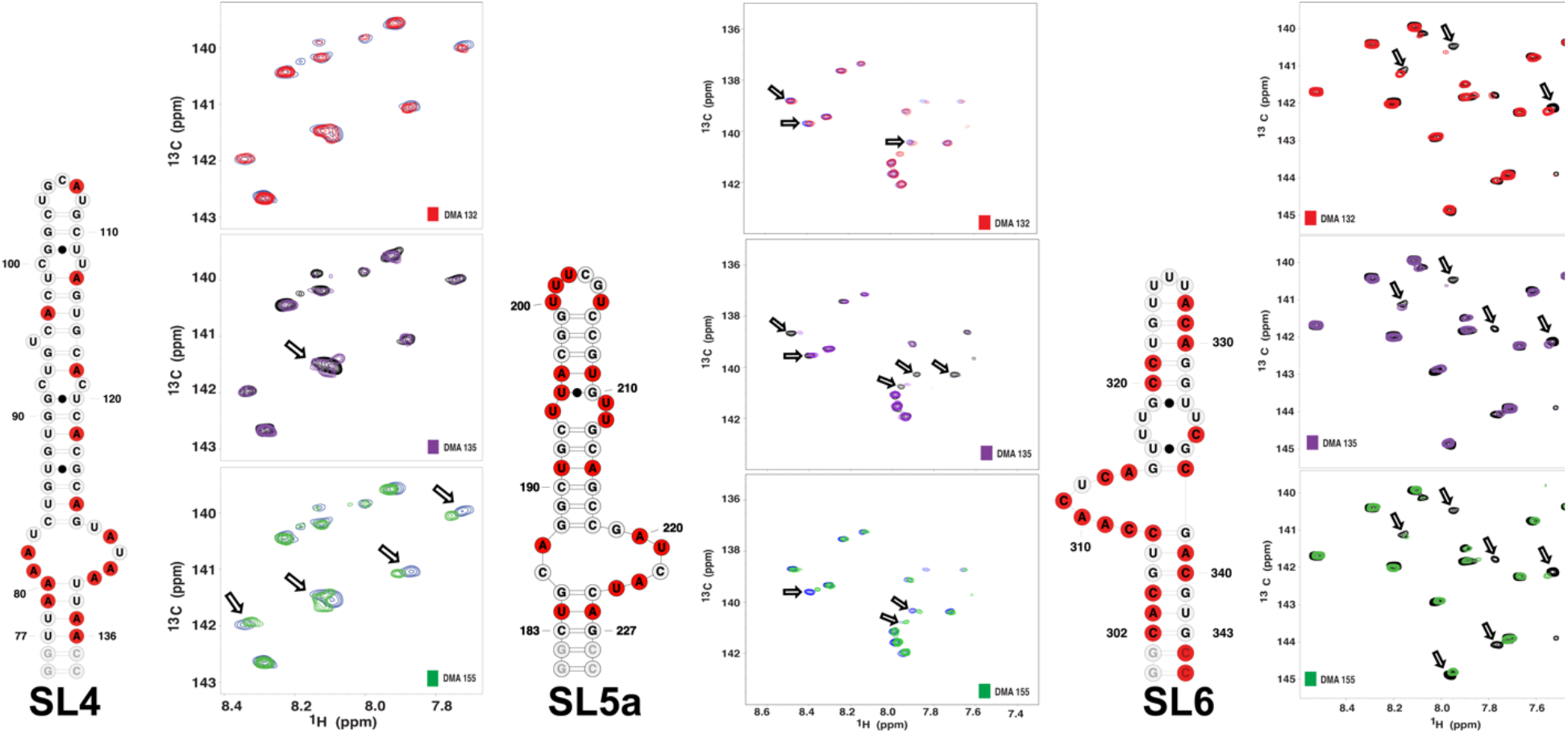
Single-point ^13^C-^1^H TROSY HSQC titrations reveal DMA-132 -135 and -155 bind with moderate affinity and specificity to SARS-CoV-2 5’-region stem loops. The spectra were recorded at 900 MHz in 100% D2O buffer of 25mM K2HPO4, 50mM KCl at pH 6.2. Temperatures (298, 303 or 308K) were optimized for each RNA construct to maximize the number of observed correlation peaks. The total RNA concentrations were set to 100 μM while titrating 5-fold excess DMA.

### VI. *In silico* ligand screening of DMA focused library against 5’-UTR and adjacent RNA structures

To generate potential 3D models of each SL, we used Fragment Assembly of RNA with Full-Atom Refinement (FARFAR). We chose FARFAR to generate preliminary models as it has consistently been demonstrated to be the most accurate RNA 3D prediction algorithm.^33^ We generated between 5,000 and 100,000 models for each SL and then generated ∼10 representative clusters.

The lowest energy conformation of each cluster was used to generate an ensemble for each SL that was submitted to ICM pocket finder to find and characterize possible binding pockets (**Table S1**). Notably, SL2, which showed no change in ^13^C -^1^H HSQC NMR chemical shifts upon small molecule addition, did not have any identifiable binding pocket. SLs 1, 3, and 5b presented binding pockets with low to intermediate scores in terms of volume, area, hydrophobicity, burriedness and druggability score (DLID) parameters often used to describe a binding pocket’s fitness. These three structures presented minor chemical shift perturbations upon small molecule addition. ^13^C-^1^H TROSY HSQC NMR experiments showed significant changes upon small molecule binding to SL4, SL5b and SL6 (**Figure 8**), all of which had the highest scores in all parameters (**Figure 9**). Most importantly, the structure that presents the highest chemical shift perturbations upon binding to the small molecule leads, SL6, is the only SL to have a binding pocket with positive DLID score (0.45, **Table S1**). DLID score has been found as a good predictor of druggability of protein binding pockets^34^ but often presents highly negative scores in RNA due to the charged backbone of RNA.^35^ Overall, this binding pocket analysis strongly correlates with NMR experimental data.

**Figure 9.**
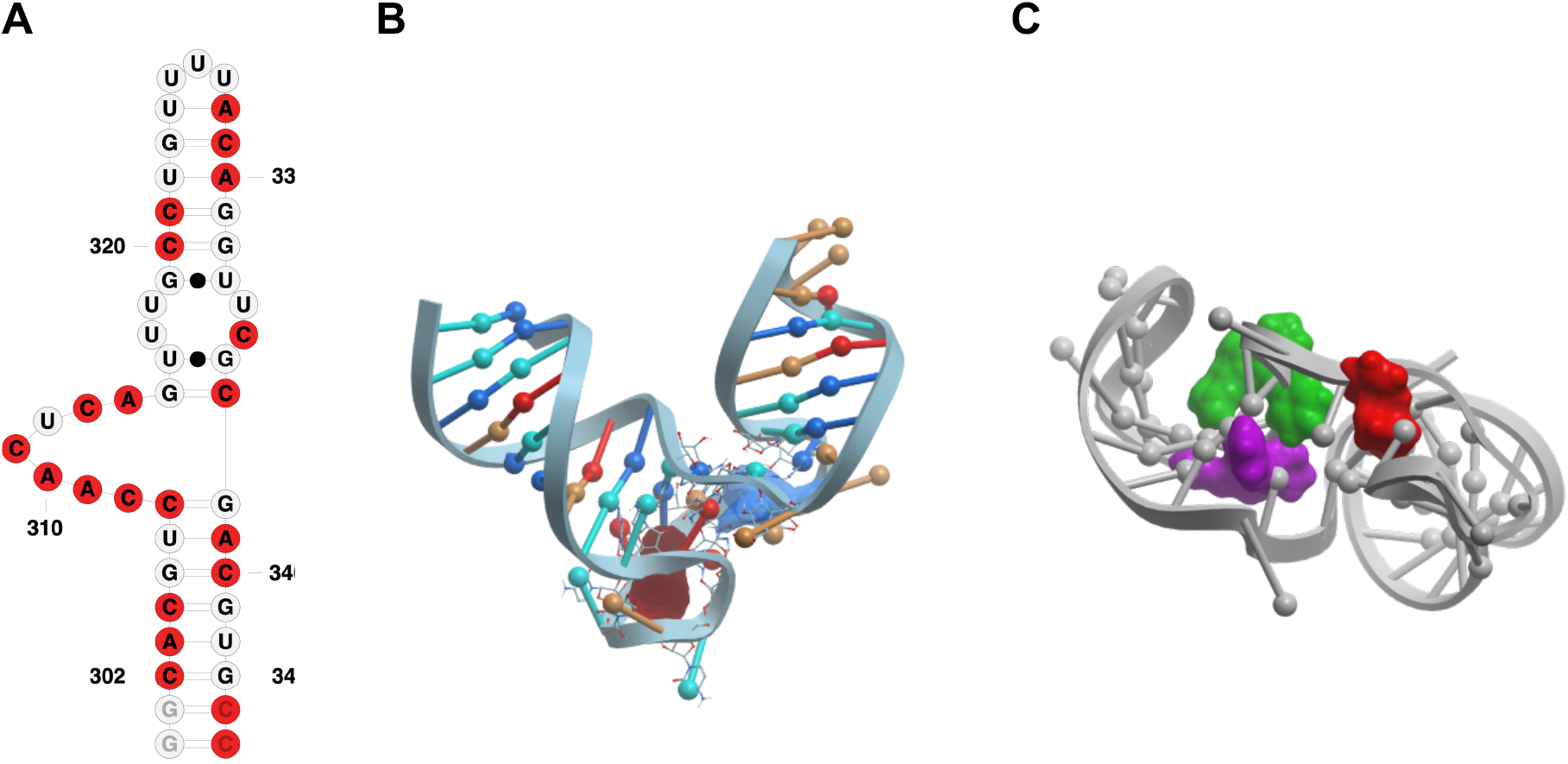
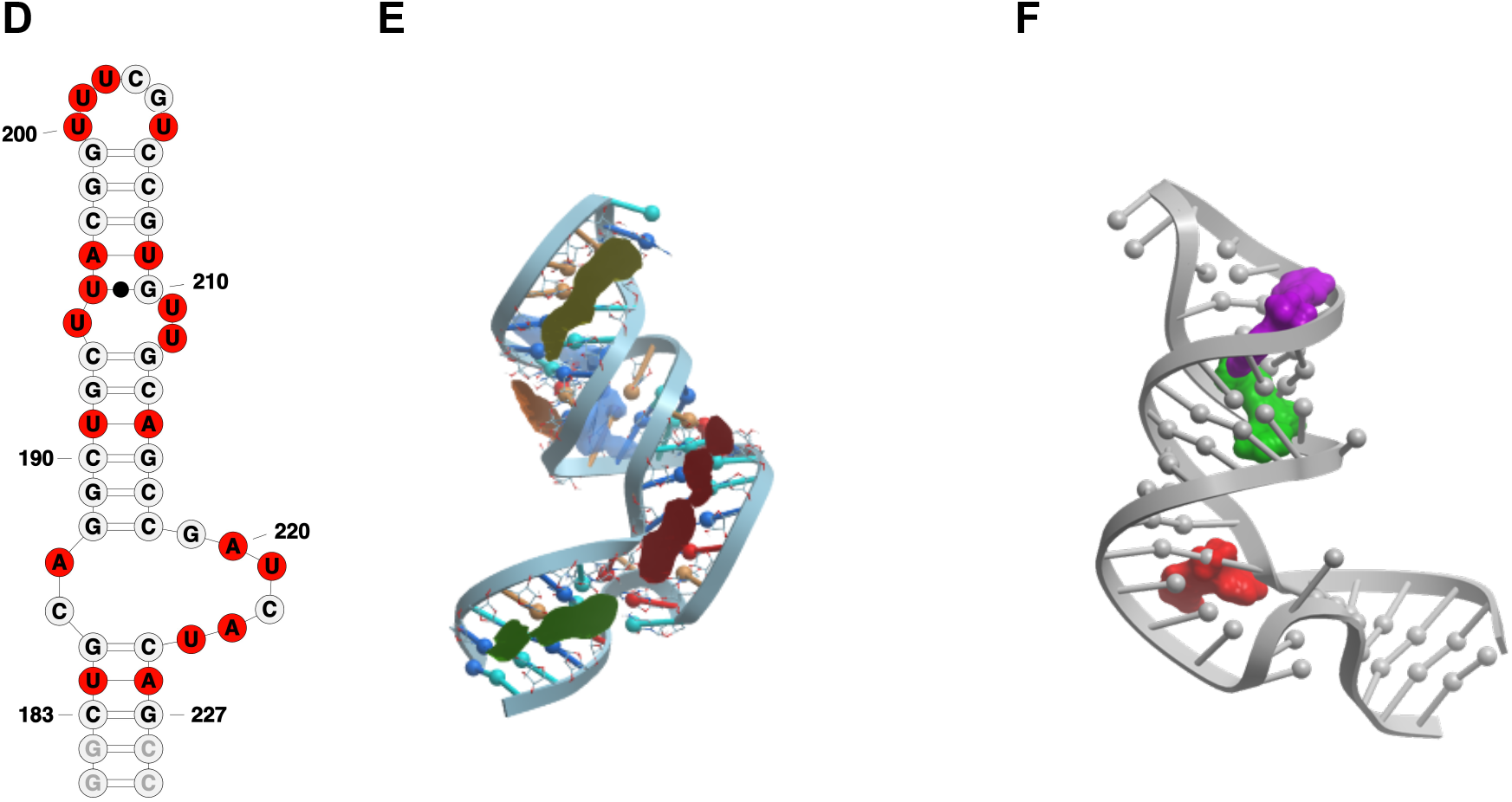
(A) Secondary structure representation of SL6 construct used in NMR studies;(B) SL6 3D model with binding pockets (red and blue) identified via ICM pocket finder; (C) Conformer that yielded the best docking scores for the three hit molecules, namely DMA 132 (red), DMA 135 (green), and DMA 155 (purple) represented in space filling model; (D) secondary structure representation of SL5a construct used in NMR studies; ; (E) SL5a 3D model with binding pockets highlighted in different colors identified via ICM pocket finder; (F) conformation and binding pockets that yielded the best docking scores of the three hit molecules DMA 132 (red); DMA 135 (green), and DMA 155 (purple).

We then docked our published 55 member DMA library against the clustered FARFAR-generated stem loop structures (**Figure S6-S11)**. In line with the results from the pocket analysis and NMR experimental data, the overall number of hits across constructs is highest for SL4, SL5a, and SL6, corroborating their potential as therapeutic targets. Hit ligands DMA-132, 135, and 155 were among the top predicted binding ligands. Furthermore, if the docked structures are limited to the sub-library of DMAs tested against OC43 and SARS-CoV-2 the three hit ligands score in the top 5 of every stem loop. Notably, when analyzing the binding location that yielded the best docking scores of the three hit molecules against SL6 we note that while DMA 135 and 155 bind best in one binding pocket (**Figure 9C** in blue), DMA 132 docks best in the adjacent binding pocket (**Figure 9C** red). When comparing the NMR profiles of the three small molecules we note that DMA 135 and 155 have a similar perturbation pattern, while DMA 132 has a slightly different profile, suggesting a possible different binding mode. This trend is recapitulated with SL5a where all three hits dock in three separate but adjacent binding pockets (**Figure 9E**,**F**) and, while all three perturb some common signals, all three molecules have a distinct NMR profile. While preliminary, this docking analysis corroborates the NMR data and the fitness of these ligands for SARS-CoV-2 5’-stem loop targeting and supports its utility as a tool in the identification of new SARS-CoV-2 RNA-targeting ligands.

## Discussion

Screening of synthetic RNA-focused libraries in recent years has provided the field with many RNA-binding bioactive small molecules and some of the highest hit rates among small molecule screening approaches.^36^ Amiloride, a known RNA-binding scaffold, has been synthetically tuned for a range of RNA secondary structures and recently yielded a novel antiviral lead for the treatment of enterovirus 71. In this case, amiloride inhibited viral translation by binding the viral 5’-UTR and modulating RNA:host protein interactions. SARS-CoV-2 also contains a highly conserved 5’-end^37^ that is reported to play a crucial role in viral replication and hijacking of host cell translational machinery. The presence of multiple bulge or internal loops, the secondary structural elements that amilorides have been reported to bind most effectively, makes the 5’-UTR and the adjacent SL6 ideal therapeutically relevant targets for small molecule probing. An initial DMA focused library screen against OC43-infected Vero E6 cells allowed for the identification of three lead compounds, namely DMA-132, -135, and -155 that significantly reduced viral titer. Initial structure-activity relationships could also be resolved, highlighting the critical substitution of the dimethylamine group at the C5 position and rigid aromatic substituents at the C6 position. These molecules were also found to be active against SARS-CoV-2 in both screening-format qRT-PCR assays and authentic antiviral titers. Luciferase assays revealed the presence of the 5’-UTR and proximal region as necessary and sufficient for translation inhibition upon small molecule treatment.

NMR profiling of leads DMA-132, -135 and -155 against each of the major 5’UTR and adjacent stem loop domains revealed that the DMAs bind preferentially to SLs that contain large internal or bulge loops. Notably SL6, which contains a moderately conserved and weakly paired ∼16-nt bulge loop, showed the most significant CSPs when titrated with DMAs. Interestingly, the 5’-side of the SL6 bulge loop has been proposed to interact with the SARS-CoV-2 Nucleocapsid (N) protein under phase separation conditions and may also impact genome packaging.^38^ We will investigate the modulation of the N protein:SL6 interaction as a potential antiviral mechanism in future work.

Finally, *in silico* analysis corroborated the experimental trend observed with NMR experiments, identifying some of the stem loops with predicted bulges (SL4, SL5a, SL6) as those with binding pockets with highest druggability. Furthermore, the small molecules that showed highest antiviral activity, namely DMA-132, -135, and -155 scored highest in the stem loops that reported significant chemical shift perturbations upon small molecule binding.

### Summary and Outlook

In summary, we herein identified drug-like small molecules that reduce SARS-CoV-2 replication and are the first antivirals to target the conserved RNA stem loops in the 5’-end region of SARS-CoV-2. Work is underway to further characterize the mode of action of these ligands, particularly putative impacts on RNA:protein interactions and specific steps in the viral replication cycle. Once characterized, we expect these amiloride-based ligands to serve as chemical biology tools to help understand CoV RNA molecular biology, such as N-dependent genome packaging and other cellular stages of the viral RNA replication process. Importantly, we have established an efficient framework to identify novel RNA-targeted CoV antivirals that will serve not only the SARS-CoV-2 pandemic but future coronavirus pandemics.

## Materials and Methods

### Analysis of sequence conservation

Representative betacoronavirus sequences were selected according to official taxonomy as represented by the International Committee for the Taxonomy of Viruses.^39^ Multiple sequence alignments of coronavirus sequences were constructed using BLAST^40^] and MAAFT^41^ as implemented in GLUE.^42^ Alignments were manually inspected and adjusted using Se-Al. Position coverage and percentage identity were calculated and visualized using JalView.^43^

### Cells and virus

Vero E6 (African green monkey kidney; ATCC CRL-1586) cells were cultured in MEM medium supplemented with 10% FBS (ThermoFisher Scientific) and maintained at 33°C. Cells were infected with human coronavirus OC43 (ATCC VR-1558) at indicated MOI (multiplicity of infection) and incubated 1 hr at 33°C for adsorption. Unbound virus was removed, and cells were refed fresh medium with various concentrations of DMAs. Media from infected cells were harvested 24 hrs post-infection, and virus titers were determined by plaque assay on Vero E6 cells. SARS-CoV-2 (USA-WA1/2020 strain; BEI Resources) was propagated and titered on VeroE6 cells, with sequence confirmation of a P2 stock to confirm stability of the viral genome.

### Effects of SARS-CoV-2 5’- and 3’-end sequence elements on luciferase reporter activity

The reporter plasmid pCoV-2-5′UTR-FLuc-3’UTR contains the SARS-CoV-2 5’UTR and adjacent coding sequences in ORF1a fused in-frame with the firefly luciferase open reading frame, followed by the SARS-CoV-2 3’UTR. For plasmid pCoV-2-5’UTR-FLuc, the Cov-2 3’UTR as replaced with vector-encoded sequence. The plasmids were kindly provided by Dr. Shin-Ru Shih (Chang-Gung University, Taiwan; manuscript submitted). Plasmid pRL, the Renilla luciferase control reporter vector, was purchased from Promega. CoV-2-5’UTR-FLuc-3’UTR, Cov-2-5’UTR-FLuc, and RLuc RNAs were in vitro synthesized from these plasmid templates using the MEGAscript T7 Transcription Kit (ThermoFisher Scientifc).

Vero E6 cells were seeded in 24-well plates. Two hundred ng of reporter RNA, 5µl of SuperFect (Qiagen), and 400µl of MEM with 10% FBS were combined and added to one well of cells. Cells were incubated at 33°C for 4 hrs and media were changed and various concentrations of DMAs were added. Two days after transfection, IRES activity was determined by measuring Renilla luciferase (RLuc) and firefly luciferase (FLuc) activities using the Dual Luciferase Reporter Assay System (Promega).

### Cytotoxicity assays

Various concentrations of DMAs were added to Vero E6 cells in culture. The cells were incubated at 33°C for 96 hrs. Cell viability was determined by MTT assay and measured at 570 nm according to the manufacturer’s instructions (EMD Millipore). All experiments were performed in triplicate. The concentration of DMAs required to reduce cell viability to 50% of the control cells was expressed as CC_50_.

### SARS-CoV-2 antiviral assays

Analysis of SARS-CoV-2 growth in cell culture supernatants was performed using a simplified RT-qPCR assay as explained before.^32^ In brief, Vero E6 cells were infected with SARS-CoV-2 at an MOI of 0.1 i.u./cell in 96-well plates, virus inoculum was removed 1 hour post-adsorption and replaced with media containing serial dilutions of the DMA compounds. 5 μL of cell culture media containing released virions were collected at 24 hpi and processed as detailed in previous studies.^32^ Viral RNA levels were quantitated by RT-qPCR using primers specific to SARS-CoV-2 N gene and a standard curve derived from in vitro synthesized RNA encoding N.

In addition, antiviral activity was tested against SARS-CoV-2 using a TCID_50_ assay, VeroE6 cells were seeded at 1 × 10^5^ per well in a 24-well plate at 37°C for 24 hours. Cells and samples were then transferred to a Biosafety Level 3 facility. Stocks of SARS-CoV-2 were diluted in DMEM/2% FBS for a solution of 20,000 pfu/mL. Growth media was aspirated from 24-well plates and replaced with 495 µL of DMEM/2% FBS containing 20,000 pfu/mL SARS-CoV-2 for an M.O.I. of 0.1. 5 µL of DMSO or compound were then immediately diluted into each well for final concentrations of 50 μM and 10 μM. Plates were incubated at 37°C for 72 hours. Media was then harvested, centrifuged at RT for 10’ at 1,500 x *g*, then used for TCID50 assay. Serial dilutions of supernatant from the treated cells were added to VeroE6 cells in 96 well plates and cells are monitored for cytopathic effect (CPE). Viral titer was calculated from the numbers of positive wells using a modified Reed and Muench method.

### Synthesis and purification of RNA stem loop constructs present at 5’-end

SL1 through 6 of the 5’-end were *in vitro* transcribed using a standard protocol,^44, 45^ from synthetic DNA templates from Integrated DNA Technologies (Coralville, IA). The 3-6 ml reactions involved the use of purified recombinant T7 RNA polymerase expressed in BL21 (DE3) cells. Depending on the labeling scheme of each RNA, double labeled, ^13^C and ^15^N, rNTPs and unlabeled rNTPs were utilized in the reaction. Nucleotide labeling of the SLs was based on the abundance of nucleotides in bulges and loops. The labeling pattern was as follows: SL1 C(^13^C,^15^N)-labeled, SL2 U(^13^C,^15^N)-labeled, SL3 AU(^13^C,^15^N)-labeled, SL4 A(^13^C,^15^N)-labeled, SL5a AU(^13^C,^15^N)-labeled, SL5b AU(^13^C,^15^N)-labeled, and SL6 AC(^13^C,^15^N)-labeled.

Next, the SLs were purified on denaturing gels, ranging from 8 to 16%, and extracted using electroelution. After desalting with a Millipore Amicon Ultra-4 centrifugal filter, the RNAs were annealed by heating for 2 min at 95 °C and flash-cooled on ice. The samples were thoroughly washed and concentrated down with a 100% D_2_O buffer of 25mM K_2_HPO_4_, 50mM KCl at pH 6.2. Using the NanoDrop™ 2000 software (Thermo Fisher), the theoretical extinction coefficients of the SLs were calculated in order to determine RNA concentrations. Samples for NMR titrations contained 100μM of RNA in D_2_O buffer with a final volume of 200μL.

### NMR profiling of DMA interactions with 5’ end structures

A 900 MHz spectrometer was used to record all NMR data. The ^1^H-^13^C HSQC titrations were recorded with 100 μM of selectively labeled SL1 through 6 in a 100% D_2_O buffer of 25mM K_2_HPO_4_, 50mM KCl at pH 6.2 with a 200μL sample volume. Titrations of the DMA molecules into the different SLs were collected at a molar ratio of 5:1, DMA to RNA, at a temperature of either 298, 303, or 308K. For each construct, temperature optimization experiments were done in order to determine the optimum temperature to conduct the titrations with. The NMR spectra were processed using NMRPipie/NMRDraw^46^ and analyzed with NMRView^47^ J or Sparky^48^

### Virtual Ligand Screening against SARS-CoV-2 5’-end RNA structures

#### FARFAR model generation

FARFAR is part of the Rosetta 3 software package can be obtained for free academic usage (https://www.rosettacommons.org/software/license-and-download). Each SL was generated through the following protocol.

*rna_helix.py* is a python wrapper for the Rosetta executable *rna_helix. rna_helix.py* is available in $ROSETTA/tools/rna_tools/bin, where $ROSETTA is the Rosetta installation path. Below are the commands to generate SL1.

*rna_helix.py -seq gguu aacc -resnum 1-4 24-27 -o helix_1.pdb*

*rna_helix.py -seq acc ggu -resnum 8-10 17-19 -o helix_2.pdb*

For each helix in a SL other than the nucleotides that flank bulges and loops we prebuild as idealized A-form helices with the above commands. Modeling helical residues as idealized A-form significantly reduces computational time allowing for more models to be built.

FARFAR modeling is performed through the *rna_denovo* executable.

*rna_denovo -nstruct 1000 -s helix_1.pdb helix_2.pdb -fasta input.fasta -secstruct_file input.secstruct -minimize_rna true -out:file:silent farfar.out*

where *-nstruct* is the max number of models requested, *-fasta* is the path of a fasta file containing the RNA sequence, *-secstruct* is the path of a file (input.secstruct) containing the RNA secondary structure in dot-bracket notation, *-minimize_rna true* minimizes the RNA after fragment assembly, *-s* specifies the path to the pdb files that contain static structures of our helices, and -out:file:silent specifies the output file path to store all generated models. Each SL was run on 100 cores for 24 hours, the number of generated models is reported in table FARFAR (Table SX).

To reduce the number of models to dock against we performed fixed-width clustering using the rna_cluster executable.

rna_cluster -in:file:silent farfar.out -nstruct 15 -cluster:radius RADIUS

Where -in:file:silent is a silent file of all models for a given SL, -nstruct is the max number of clusters requested, and -cluster:radius is max distance in heavy-atom RMSD between members of the same cluster.

#### Small molecule docking

Virtual docking simulations were performed using ICM^49^ (Molsoft LLC. La Jolla, CA). employing the SARS-CoV-2 5’-UTR structures obtained from FARFAR modeling and clustering. RNA structural elements’ binding pockets were defined using ICM Pocket Finder module and all the small molecules protonation states were adjusted to pH=7.0 using ChemAxon^**©**^ (www.chemaxon.com). RNA ensembles were then combined into conformational stacks using ICM’s “impose conformations.” Then, to create “flexible receptors” to dock against that would reflect all of the conformations of each structure, ICM’s “create 4D grid” function was used for each docking project. Each of the structures was then docked against a library of 55 DMA molecules. The DMA library was saved in .sdf file format, which was indexed for VLS using ICM-Pro. The virtual screening simulation was implemented with a conformational search and optimization with a limit of 10 conformers per molecule. The thoroughness was left at level 10.

## Acknowledgements

The authors are grateful for intellectual discussions and advice from Alice Telisnitsky, Silvi Rouskin, Nathan Sherer, and Christopher Laudeman. We thank Neeraj Patwardhan and Aline Umuhire Juru for contributing DMA molecules included in the initial screen.

The authors acknowledge funding from the following sources:

NIGMS R35GM124785 ; Duke University to A.H.

NIGMS, GM126833 to G.B. and B.T.

Tobacco Settlement Fund 21-5734-0010 to J.Y.

Medical Research Council of the United Kingdom (MC_UU_12014/12) to R.G.

## Conflicts of Interest

The authors declare no conflicts of interest.

## Notes

### Competing Interest Statement

The authors have declared no competing interest.

