## Supplemental Information for "Amilorides inhibit SARS-CoV-2 replication in vitro by targeting RNA structures"

† Co-first authors

‡ Co-second authors

* Corresponding authors

Contact:

### Table of Contents

*I. Small Molecule screening against OC43 with Vero E6 cells* ______________ *S2*

*II. Replicates of Antiviral effects of lead DMAs* ______________ *S2*

*III. Small molecule toxicity in Vero E6 cells* ______________ *S3*

*IV. Q-RT-PCR assay for dose dependent antiviral activity* ______________ *S3*

*V. $^{13}$C -HSQC NMR experiments SARS-CoV-2 5'-end RNA structures with DMAs* ______________ *S4*

*VI. In silico screening of DMA focused library against 5'-end RNA structures* ______________ *S5-S7*

### I. Small Molecule screening against OC43 with Vero E6 cells

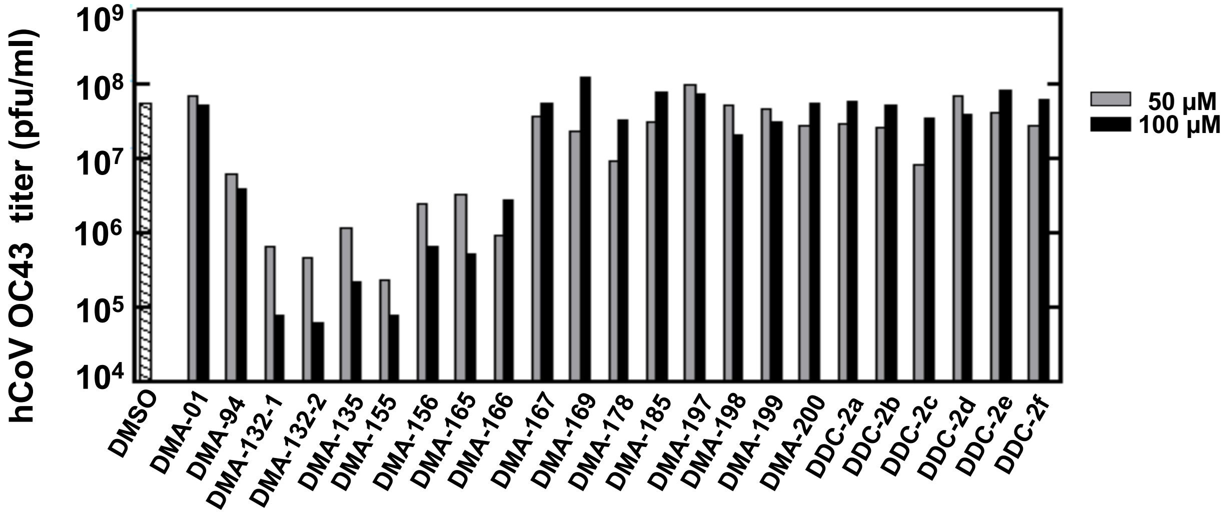


**Figure S1**. Vero E6 cells were infected with human coronavirus OC43 at an MOI 1. Various concentrations of DMAs were added to the cells. Media were harvested 24 hr post-infection and assayed for infectious virus by plaque formation with Vero E6 cells.

### II. Replicates of antiviral effects of lead DMAs

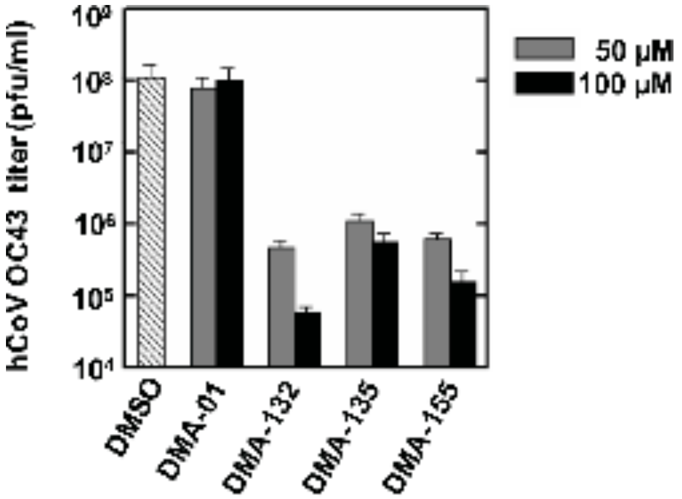


**Figure S2**. Vero E6 cells were infected with human coronavirus OC43 at an MOI 1. DMAs were added to the cells at 50 µM or 100 µM. Media were harvested 24 hr post-infection and assayed for infectious virus by plaque formation with Vero E6 cells. Mean values and standard deviations from three independent experiments are shown in the bar graphs. $P < 0.01$ for all three DMAs.

### III. Small molecule toxicity in Vero E6 cells

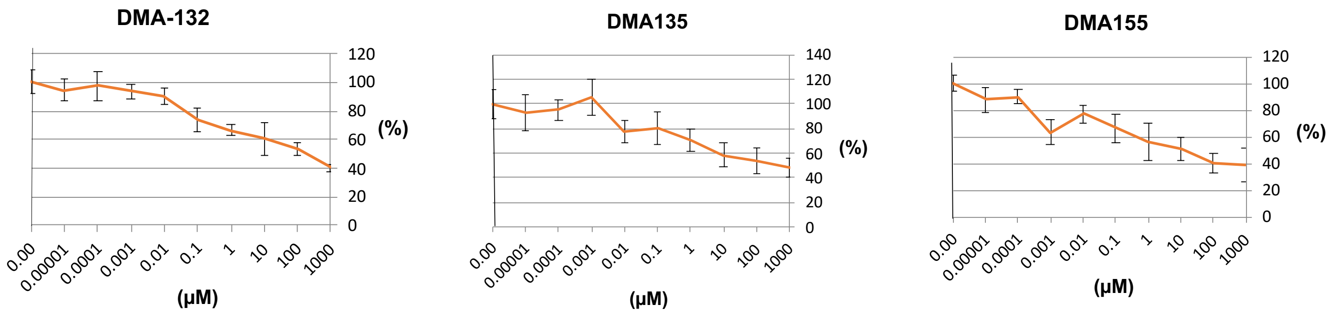


**Figure S3**. Cytotoxicity assay shows the $CC_{50}$ of DMA-132 and -135 in Vero E6 cells were > 100 μM. $CC_{50}$ of DMA-155 was about 90 μM. Various concentrations of DMAs were added to Vero E6 cells. Cells were incubated at 33°C for 96 hrs. Cell viability was determined by MTT assay with measurements at 570 nm according to the manufacturer's instructions (EMD Millipore). All experiments were performed in triplicate.

### IV. Q-RT-PCR assay for dose dependent antiviral activity

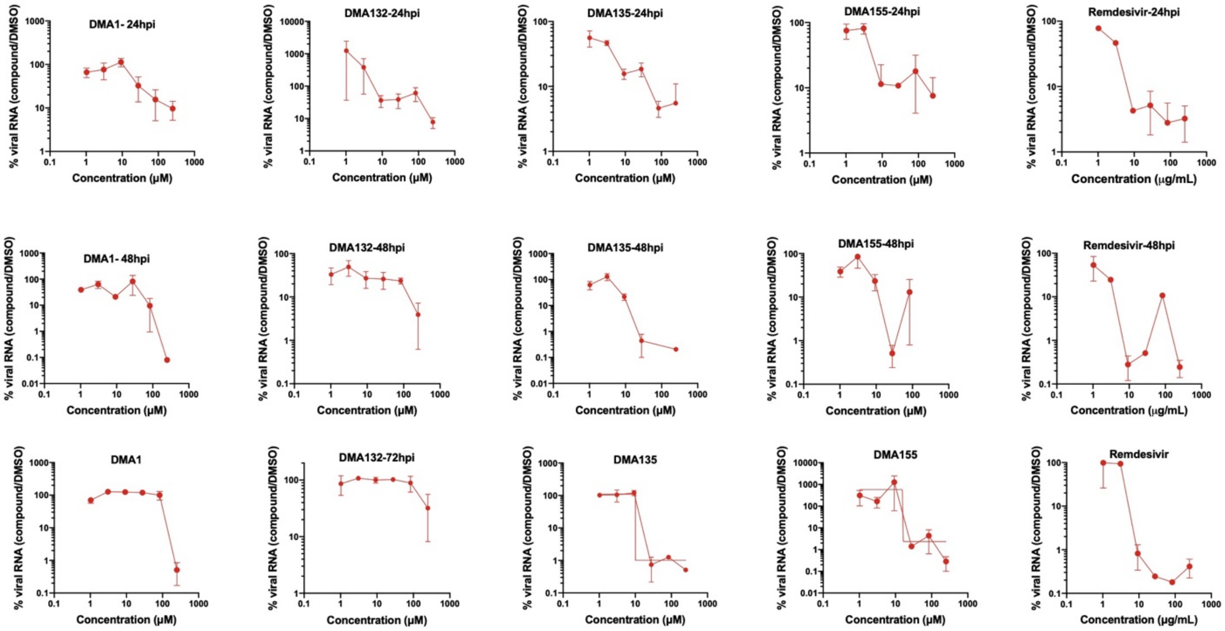


**Figure S4**. Vero E6 cells were infected with SARS-CoV-2 at MOI: 0.1 i.u./cell and the indicated DMA compounds were added on cells following virus adsorption for 1 hr. Cell culture supernatants were collected and analyzed by a Q-RT-PCR assay using N-specific primers. Data show the relative percentage of viral RNA in DMA-treated samples compared to mock-treated samples at 24, 48, and 72 hours. Data are from two independent experiments and error bars show the range.

### V. $^{13}$C -HSQC NMR experiments SARS-CoV2 5’-end RNA structures with DMAs

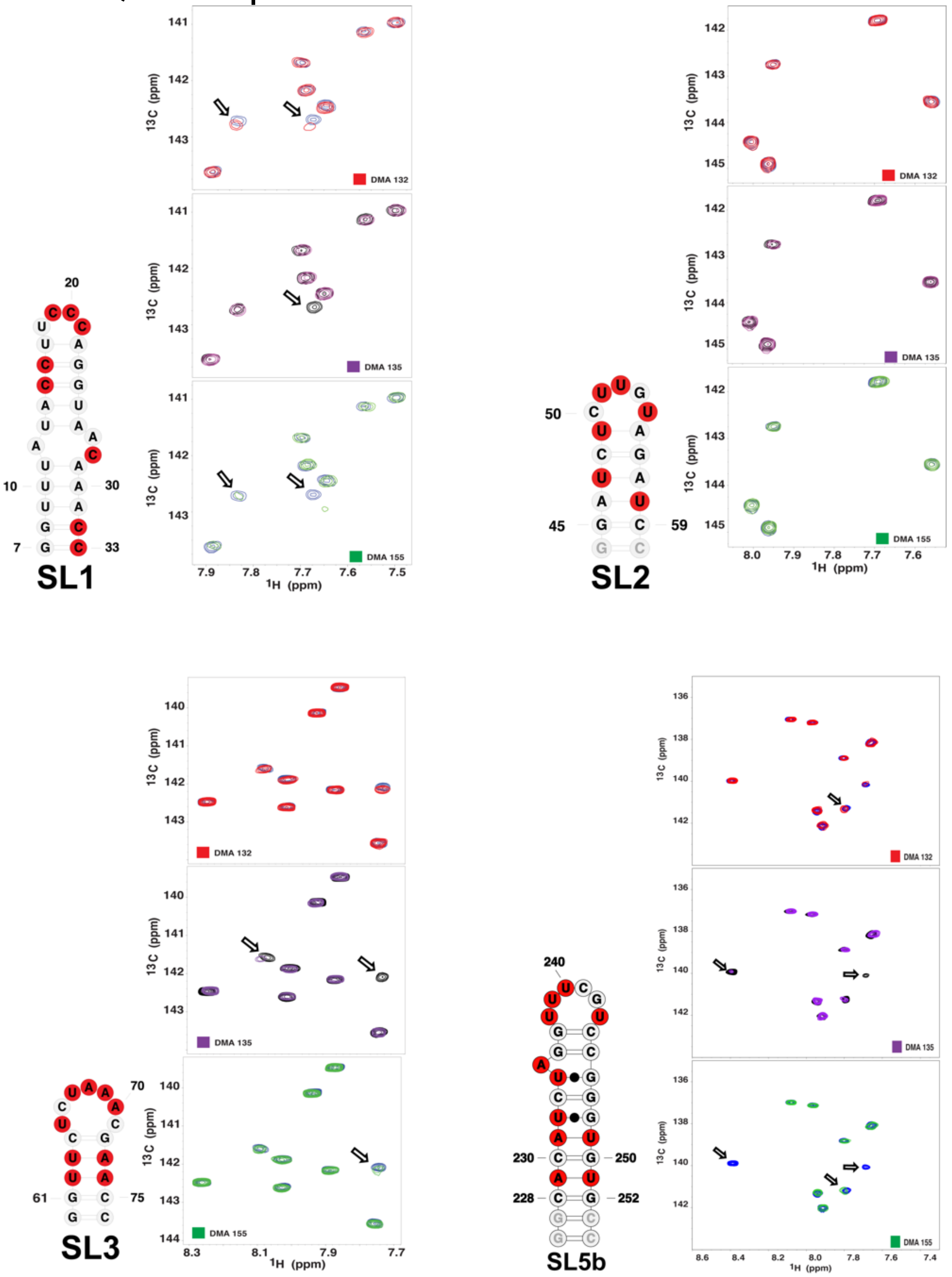


**Figure S5.** Single-point $^{13}$C-$^{1}$H TROSY HSQC titrations reveal DMA-132 -135 and -155 bind with moderate affinity and specificity to SARS-CoV-2 5’-region stem loops. The spectra were recorded at 900 MHz in 100% $D_2O$ buffer of 25mM $K_2HPO_4$, 50mM KCl at pH 6.2. Temperatures (298, 303 or 308K) were optimized for each RNA construct to maximize the number of observed correlation peaks. The total RNA concentrations were set to 100 μM while titrating 5-fold excess DMA.

### V. *In silico* screening of DMA focused library against 5'-end RNA structures

**Table S1.** Binding pockets identified with ICM pocket finder and characterization of each binding pocket.

| SL | Volume | Area | Hydrophobicity | Buriedness | Aromatic | DLID | Radius | Nonsphericity |
|---|---|---|---|---|---|---|---|---|
| **1.4** | 132.87 | 143.87 | 0.47 | 0.86 | 0.05 | -0.63 | 3.17 | 1.14 |
| **1.2** | 111.15 | 211.04 | 0.26 | 0.56 | 0.00 | -2.42 | 2.98 | 1.89 |
| **1.1** | 108.75 | 234.70 | 0.20 | 0.56 | 0.04 | -2.59 | 2.96 | 2.13 |
| **3** | 111.58 | 124.56 | 0.52 | 0.85 | 0.00 | -0.72 | 2.99 | 1.11 |
| **4.1** | 583.35 | 615.91 | 0.37 | 0.80 | 0.09 | -0.02 | 5.18 | 1.82 |
| **4.2** | 142.86 | 192.83 | 0.35 | 0.68 | 0.02 | -1.55 | 3.24 | 1.46 |
| **4.3** | 126.79 | 168.48 | 0.40 | 0.78 | 0.08 | -1.15 | 3.12 | 1.38 |
| **5a.1** | 273.58 | 474.91 | 0.37 | 0.74 | 0.03 | -0.81 | 4.03 | 2.33 |
| **5a.2** | 174.42 | 274.74 | 0.37 | 0.70 | 0.01 | -1.29 | 3.47 | 1.82 |
| **5a.3** | 160.57 | 228.90 | 0.38 | 0.69 | 0.01 | -1.37 | 3.37 | 1.60 |
| **5a.4** | 133.51 | 206.26 | 0.31 | 0.64 | 0.00 | -1.86 | 3.17 | 1.63 |
| **5a.5** | 132.20 | 180.61 | 0.42 | 0.70 | 0.06 | -1.39 | 3.16 | 1.44 |
| **5b** | 285.30 | 310.08 | 0.49 | 0.85 | 0.14 | -0.04 | 4.08 | 1.48 |
| **6.2** | 235.09 | 215.04 | 0.61 | 0.95 | 0.11 | 0.45 | 3.83 | 1.17 |
| **6.1** | 219.49 | 239.54 | 0.57 | 0.76 | 0.20 | -0.41 | 3.74 | 1.36 |

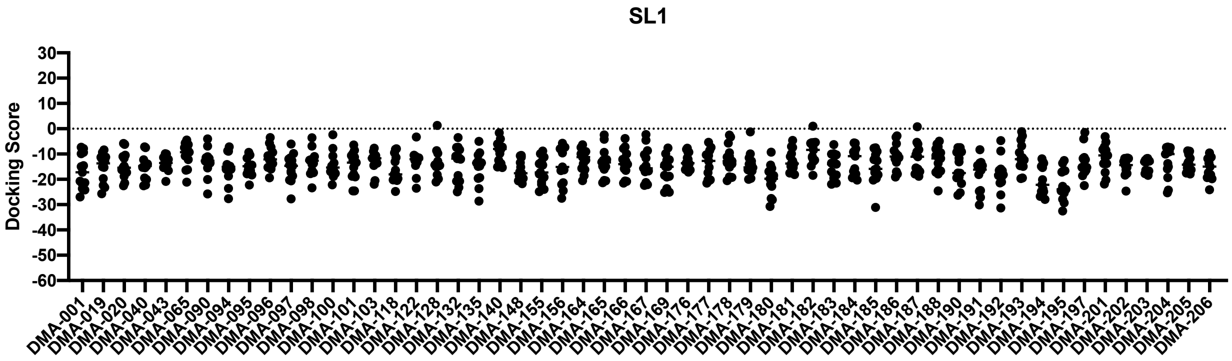


**Figure S6**. Docking score of the 55 member DMA library against stem loop 1 (SL1). The dots represent the best docking scores for each SL1 conformer present in the cluster.

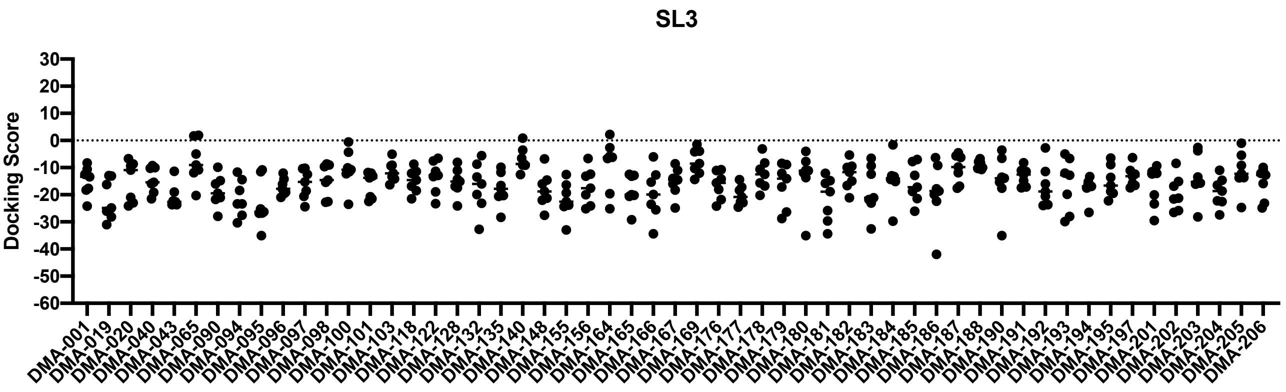


**Figure S7**. Docking score of the 55 member DMA library against stem loop 3 (SL3). The dots represent the best docking scores for each SL3 conformer present in the cluster.

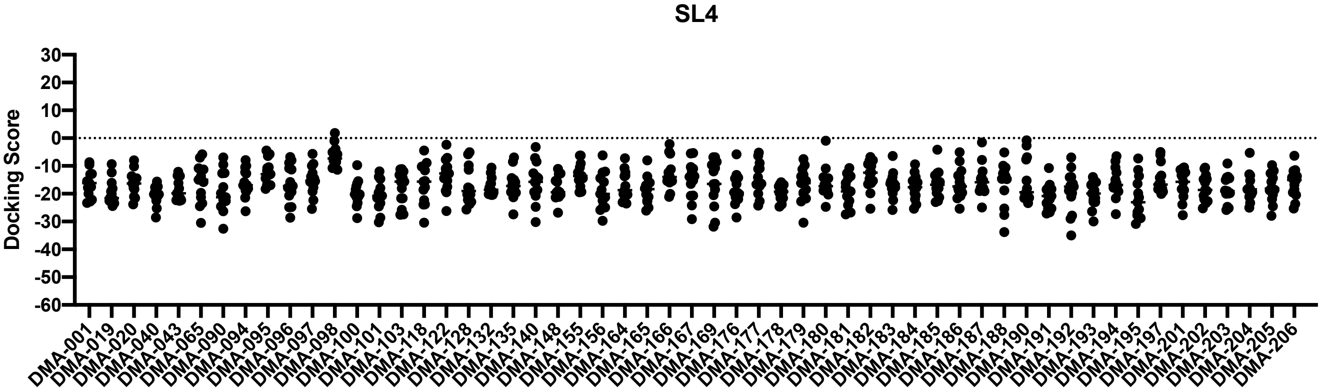


**Figure S8**. Docking score of the 55 member DMA library against stem loop 4 (SL4). The dots represent the best docking scores for each SL4 conformer present in the cluster.

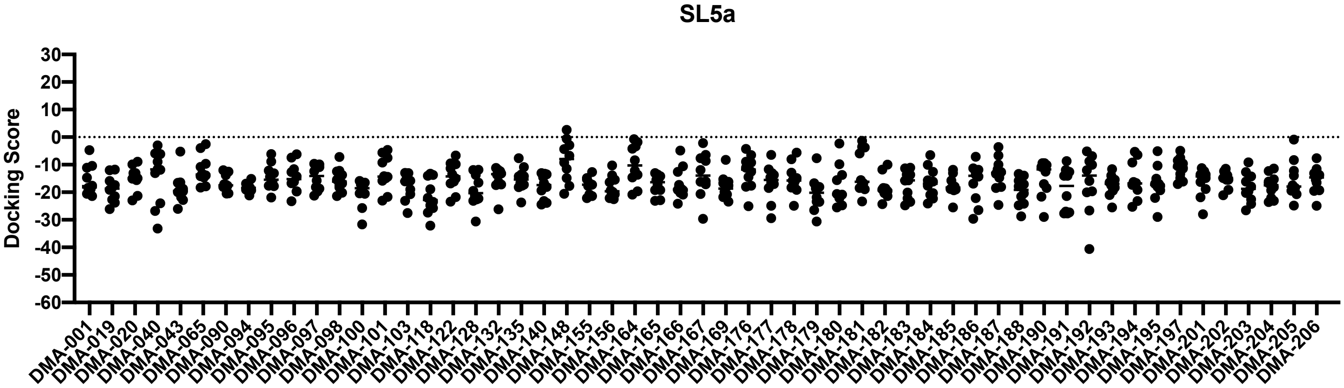


**Figure S9**. Docking score of the 55 member DMA library against stem loop 5a (SL5a). The dots represent the best docking scores for each SL5a conformer present in the cluster.

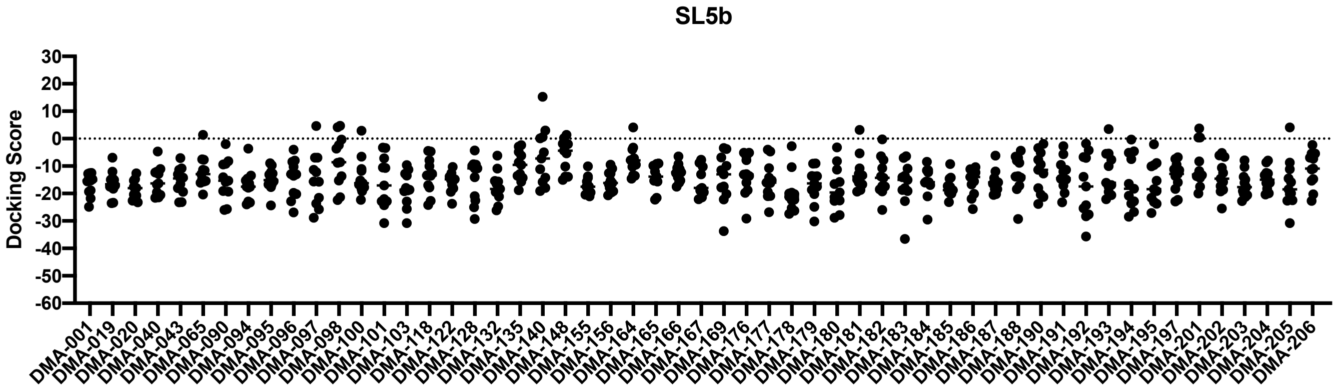


**Figure S10**. Docking score of the 55 member DMA library against stem loop 5b (SL5b). The dots represent the best docking scores for each SL5b conformer present in the cluster.

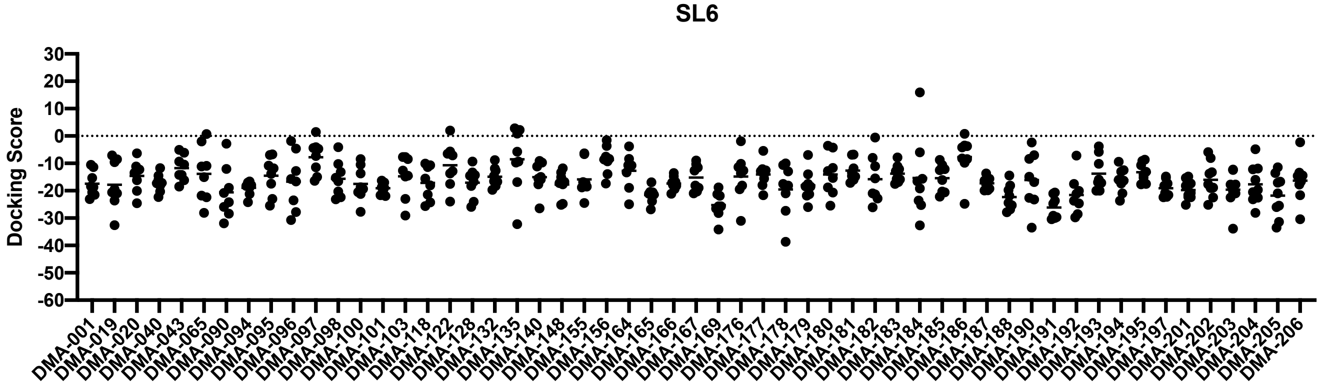


**Figure S11**. Docking score of the 55 member DMA library against stem loop 6 (SL6). The dots represent the best docking scores for each SL6 conformer present in the cluster.
